# Isolation and Genomics of multidrug-resistant *Brevundimonas diminuta* collected from patients with pertussis-like symptoms

**DOI:** 10.1101/2021.08.23.457371

**Authors:** Azadeh Safarchi, Vajihe Sadat Nikbin, Samaneh Saedi, Mojdeh Dinarvand, Mahdi Sedaghatpour, Masoumeh Nakhost Lotfi, Chin Yen Tay, Binit Lamichhane, Fereshteh Shahcheraghi

## Abstract

*Brevundinomas diminuta* is known as an opportunistic pathogen and is rarely associated with invasive infections in humans causing infection in the different parts of the body. In this study, we identified three *B. diminuta* isolates from three patients with an initial diagnosis of pertussis. Isolates were confirmed using biochemical tests and 16s rRNA sequencing. All isolates were resistant to three different classes of antimicrobial drugs including ceftazidime and ciprofloxacin. We performed Illumina whole-genome sequencing for two isolates. The results showed an average genome size of 3.25 Mbp with the G+C content 67% with multiple predicted virulence factors and genes leading to antibiotic resistance. We found an open pan-genome with 1502 core genes by analysing 13 available global *B. diminuta* isolated from different environments such as water, soil, or gentamicin fermentation residue. In the phylogenetic analysis, our isolates were grouped with *B. diminuta* isolate collected from an oral cavity of a patient in the USA.

## 1 Introduction

Brevundimonas species is a non-fermenting Gram-negative bacillus belonging to the Alphaproteobacteria class and *Caulobacteraceae* family with a G + C content of 65% to 68%. It was previously assigned to the genus *Pseudomonas* and *Caulobacter* as *Pseudomonas diminuta* [1]. However, whole-cell protein patterns, fatty acid compositions, phenotypic characteristics, and DNA base rations classified it into a new genus that currently contains 25 species mostly isolated from diverse soil and aquatic environments [2]. They are usually used as test organisms for several purposes from validating reverse-osmosis filtration devices to mitigating the toxic effects of heavy metals on plant growth in contaminated soils [3, 4]. *Brevundimonas vesicularis* and *Brevundimonas diminuta* have also been isolated from clinical environments as opportunistic pathogens from patients with cancer cystic fibrosis and, tuberculosis [5, 6, 7]. Reports also showed some sporadic incidences caused by *B. diminuta* as an emerging opportunistic pathogen causing infection in the bloodstream, intravascular catheter, urinary tract, and pleural space [4].

While *B. diminuta* is rarely reported in invasive respiratory tract infections, it has been isolated from patients with over two weeks of pleuritis symptoms such as intermittent cough, sore throat, breath-holding, and fever [8]. Here, we report the isolation of three *B. diminuta* strains from patients with pertussis-like symptoms and investigated their antimicrobial resistance profiles. We further analysed genomic features of two *B. diminuta* using whole-genome sequencing and compared them with available sequenced global *B. diminuta* isolates.

## 2 Methods and materials

### 2.1 Bacterial strains and identifications

Clinical isolates were isolated from nasal swabs of patients with acute respiratory symptoms that were sent to the bacterial reference laboratory in Pasteur Institute of Iran to confirm pertussis diagnosis. Samples were cultured on Reagan-Lowe agar supplemented with 10% defibrinated sheep blood incubated at 35-37°C for one to three days. Gram-staining, colony morphology, as well as the presence of cytochrome C, were investigated. Further biochemical characterization was performed using Analytical Profile Index (API) 20E test (BioMerieux, Inc., Hazelwood, MO).

The 16s ribosomal RNA was amplified from the extracted genome. Briefly, DNA was extracted using in-house method from pure culture and subjected to PCR amplification for a 1500 bp fragment of the 16s rRNA gene. A set of universal bacterial primers, 27F, and 1429R were used for PCR amplification. The amplicons were sequenced (Macrogen Co, South Korea) and sequence analysed using MEGA (V6)[9].

### 2.2 DNA preparation and DNA sequencing

Two isolates, IR18 and IR165, isolated in 2009 and, 2016 respectively, were selected for whole-genome sequencing. Genomic DNA was extracted from bacterial culture using a phenol-chloroform extraction method and ethanol precipitation as described previously [10]. DNA libraries were prepared with the insert size of 150 bp paired-end using NexteraXT DNA kit (Illumina) and sequenced on the Nextseq (Illumina) with a minimum coverage of 150-fold. The raw reads were submitted to the GeneBank database under BioProject No. PRJNA689668.

### 2.3 Bioinformatics and genome analysis

*de novo* assembly was performed for all sequencing data as described previously to combine reads into contigs using SPAdes (v3.7.2) [11, 12]. To determine the closest strain, the contigs were uploaded for BLAST search in the NCBI database. RAST (Rapid Annotations using Subsystem Technology and NCBI prokaryotic genome annotation pipeline (PGAP) were applied to annotate genomes and predict genes [13] [14].

Reads were also mapped against *B. diminuta* strain ATCC11568 (Taxonomy ID 751586) as a reference genome and using Burrows-Wheeler Alignment (BWA) tools (version 0.7.12) and SAMtools (v 0.1.19). We also mapped reads against *B. diminuta* strains BZC3 as the closest isolate based on BLAST results [10, 15, 16, 17].

The virulence factors were investigated using the nucleotide BLAST method against the virulence factor database [18] with an E-value <1.0 E10-6 and the results were filtered with an identity cut-off 75% and alignment length >200 bps.

Assemblies were submitted to CRISPR finder and Phaster for CRISPR (clustered interspaced short palindromic repeats) and prophage prediction in the genomes, respectively [19, 20].

Assemblies of publicly available *B. diminuta* isolates from different environmental sources (11 isolates) were downloaded from the NCBI database (Table-3) for pan-genome analysis using Prokka and Roary [21, 22]. Briefly, draft genomes were annotated by Prokka (v1.13.3) to produce gff files which were further analysed using Roary (V3.12.0) with protein BLAST using 90% identity cut-off.

### 2.4 Antibiotic resistance gene identification and susceptibility testing

The antibiotic susceptibility test was performed with broad-spectrum antibiotics such as macrolides, bata-lactam, aminoglycosides, cephalosporines, and fluoroquinolone using the disk diffusion method according to the CLSI standard guideline (CLSI, 2010) as described previously [23].

The contigs of sequenced isolates, IR18 and IR165, were investigated to find any resistance gene using genome annotation and the nucleotide BLAST method against the Comprehensive Antibiotic Resistance Databases (CARD) using 70% identity cut-off [24].

### 2.5 Phylogenomic analysis

Phylogenetic analysis using MEGA (version 7) was conducted [9]. The maximum parsimony was applied based on Tree-Bisection-Reconnection (TBR) analysis to find out the relationship between Iranian and global isolates using core genes. Bootstrap analysis was based on 500 replicates.

## 3. Results

### 3.1 Bacterial identification

Three *B. diminuta* isolates were identified during 2009-2016 from male patients with respiratory tract infections (Table-1) of which two were infants under six-month-old and one was five-years-old children from Khorasan Razavi, an Eastern province, and Mazandaran, a Northern Province, of Iran. All patients had respiratory infection symptoms including fever, vomiting, and acute cough that were initially diagnosed as whooping cough. The nasal swabs were collected and sent to the Reference Pertussis Laboratory of Pasture Institute of Iran for laboratory confirmation based on the National Pertussis guideline [25]. Bacterial isolation from nasal swabs was carried out as previously described [23, 26]. Isolates were initially confirmed as *B. pertussis* due to their growth on Bordet Gengou agar and some biochemical analysis such as catalase, oxidase and urease test. However, the molecular determination by real-time PCR targeting IS*1002*, IS*481* and the *ptx* promoter as a gold molecular standard for *B. pertussis* confirmation [26] for these three isolates was negative. Therefore, additional biochemical tests such as oxidase production and antibody neutralizing teste as well as 16s rRNA sequencing were performed. These tests suggest that all three isolates were *B. diminuta*.

**Table-1:** Clinical and biochemical characterization of *B. diminuta* isolates collected during 2009-2016 in Iran.

| Sample<br>(Isolates ID) | Patient age | Source | Symptoms | Province of<br>isolation | Year |
| --- | --- | --- | --- | --- | --- |
| IR165 | 5 years (Male) | Nasopharynx | Fever, Cough, Vomit | Mazandaran | 2016 |
| IR11 | 26 days (Male) | Nasopharynx | Fever, Cough, Vomit | Khorasan Razavi | 2009 |
| IR18 | 4 months (Male) | Nasopharynx | Cough, Vomit | Khorasan Razavi | 2009 |

### 3.2 General genome features

Two *B. diminuta* isolates, IR18 and IR165, with six years interval of collection were selected for whole-genome sequencing to understand their genomic features. IR18 was isolated in 2010 and IR165 isolated in 2016 from 1.5-month-old infant and five-years-old children, respectively. Our results revealed 50 contigs for each assembled fasta file with the average G+C% content of 67% and genome size 3161756 for IR18 and 3341181 for IR165 which is similar to the average genome size 3.3 Mbp reported for this strain [6].

The annotation by RAST and prokaryotic genome annotation pipeline (PGAP) showed 3077 and 3285 potential protein sequences (CDSs) for IR18 and IR165 respectively (Supplementary File-1), which is consistent with the average number of proteins sequences of available global isolates. Genome annotation showed that the majority of the genes belonged to functional groups involved in cell synthesis and metabolisms (Fig. 1) Three rRNAsincluding one large and one small subunit of ribosome, and one 5sRNA were identified in the genomes. Furthermore, 48 and 49 tRNA found in IR18 and IR165, respectively.

**Figure-1.**
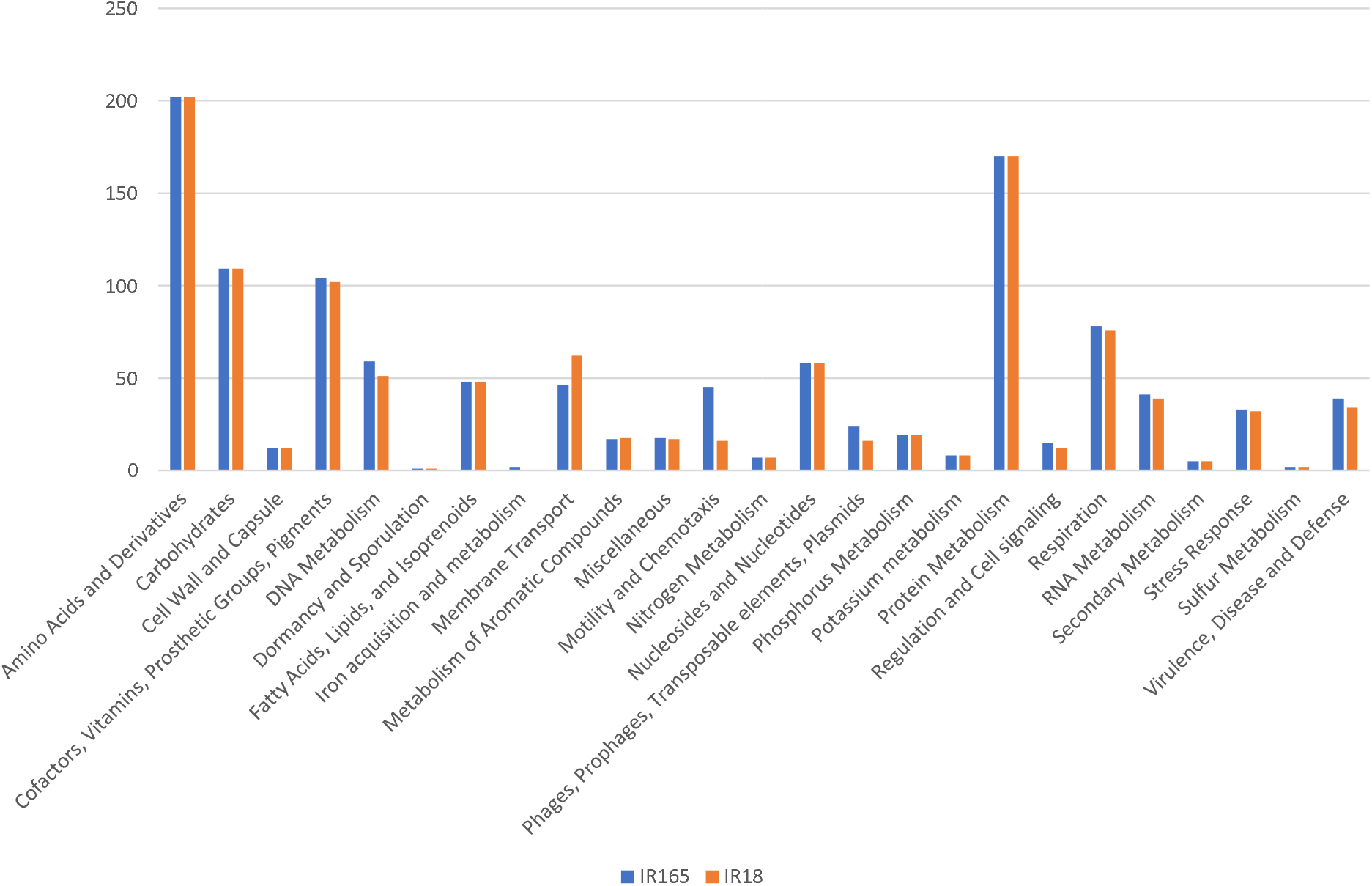
Subsystem information of *B. diminuta* IR165 and IR18 annotated by RAST.

Only 65.6% and 55.4% of reads in IR18 and IR165 respectively were mapped to *B. diminuta* ATCC11568 (Taxonomy ID 751586) while 98.16% and 74.9% of the reads can be mapped to *B. diminuta* strain BZC3 for IR18 and IR165, respectively. *B. diminuta* strain BZC3 was isolated from gentamicin fermentation residue in China with high gentamicin degradation efficiency [27].

### 3.3 Genome-based identification of virulence factors

The putative virulence factors in *B. diminuta* IR165 and IR18 were predicted by genome annotation using NCBI RAST as well as BLAST searches against the virulence factor database (VFDB). Various virulence factors were identified mostly categorised as secretion systems, adhesion factors, motility, iron uptake and toxin (Supplementary file 2). Putative genes encoding proteins involved in type II and III secretion systems were found in both isolates. The gene, *ompA*, encoding OmpA outer membrane family protein was identified in both genomes as adhesion protein. In the motility category, genes that encode flagella (Fli family proteins MotB) were detected in both isolates and may play an important role in adhesion, virulence factor secretion and biofilm formations in addition to motility. In the toxin-antitoxin category, we identified genes encoding type II toxin antitoxin (TA) family, YeoB-YefM, which have been reported recently to be involved in antibiotic resistance, biofilm formation and host infection as well as environmental stresses [28, 29]. Another TA-related protein identified in the genomes is a *ratA* gene encoding Ribosome association toxin A, RatA, that reported to effectively block the translation initiation step in *E*.*coli* [30]. Genes encoding chemotaxis proteins (*cheB, cheD, cheR, cheY and cheYIII*) were also identified in both genomes and may coordinate the flagellar movement in bacteria to survive better in different environment [31]. We also found various genes involving in the iron uptake including *iscR, sufA, sufB, sufC, sufD, sufD, fieF, fecD* in both genomes as well as hemolysin-related proteins.

### 3.4 Mobile elements

Phaster-tools was used to identify phage region in our genomes. It categorizes the potential prophage sequences in the genomes as intact, incomplete, or questionable [20]. One intact and one incomplete phage region were predicted to be present in each of our genomes. IR18 contained one intact region, PHAGE_Pseudo_phiCTX_NC_003278(9), which consists of 22.9 kb and a total of 30 CDS and GC content 67.35%. The predicted phage in IR18 shared sequence similarity with *Pseudomonas* species phage comprising 35 kb length and 42 protein-coding sequences [32]. IR18, also contained one incomplete prophage region with shared sequence similarity to PHAGE_Faecal_FP_Brigit_NC_047909(1) with a total of 11 protein-coding sequences and 7.4 kb length and G+C content of 63.93%.

IR165 had also one intact phage region with a total CDS 32 and GC content 67.35%. It also harbors an incomplete prophage region that consists of 42.4 kb and 37 protein-coding sequences with GC content of 62%.

We also investigated the presence of clustered interspaced short palindromic repeats (CRISPR) in the genomes using CRISPR finder tool [19]. CIRSPR-cas system is composed of CRISPR sequence that codes for a CRISPR RNA (crRNA) and cas gene cassettes that codes for CRISPR-associated (cas) proteins [33]. We observed one CRISR/cas system for IR18 and two for IR165 (Supplementary file 3). The candidate cas protein sequences were identified as helicase protein family such as DEAD-box helicases. DEAD-box helicases involve in unwinding nucleic acid and various aspects of RNA metabolism, including nuclear transcription, pre mRNA splicing, ribosome biogenesis, nucleocytoplasmic transport, translation, RNA decay and organellar gene expression as well as antiviral defence [34]. These proteins are essential for phage defence and classified as CRISPR-associated gene3 (Cas3) family in the annotation of CRISPR/cas system [35, 36].

### 3.5 Antimicrobial drug resistance mechanisms

We performed a disk diffusion method to investigate the bacterial resistance to broad-spectrum antimicrobials of various antibiotics. All three *B. diminuta* isolates were susceptible to azithromycin, gentamicin, imipenem, amikacin, cefotaxime and sulfamethoxazole/trimethoprim (Table-2). They showed different levels of resistance to ceftazidime and ceftriaxone from the cephalosporines family due to the presence of *blaVIM-2, blaVIM-13* and other beta-lactamase genes encoding metallo-β-lactamase proteins that hydrolyse cephalosporins (supplementary file-2). Furthermore, they showed a high level of resistance to ciprofloxacin and intermediate resistance to levofloxacin.

**Table-2:**
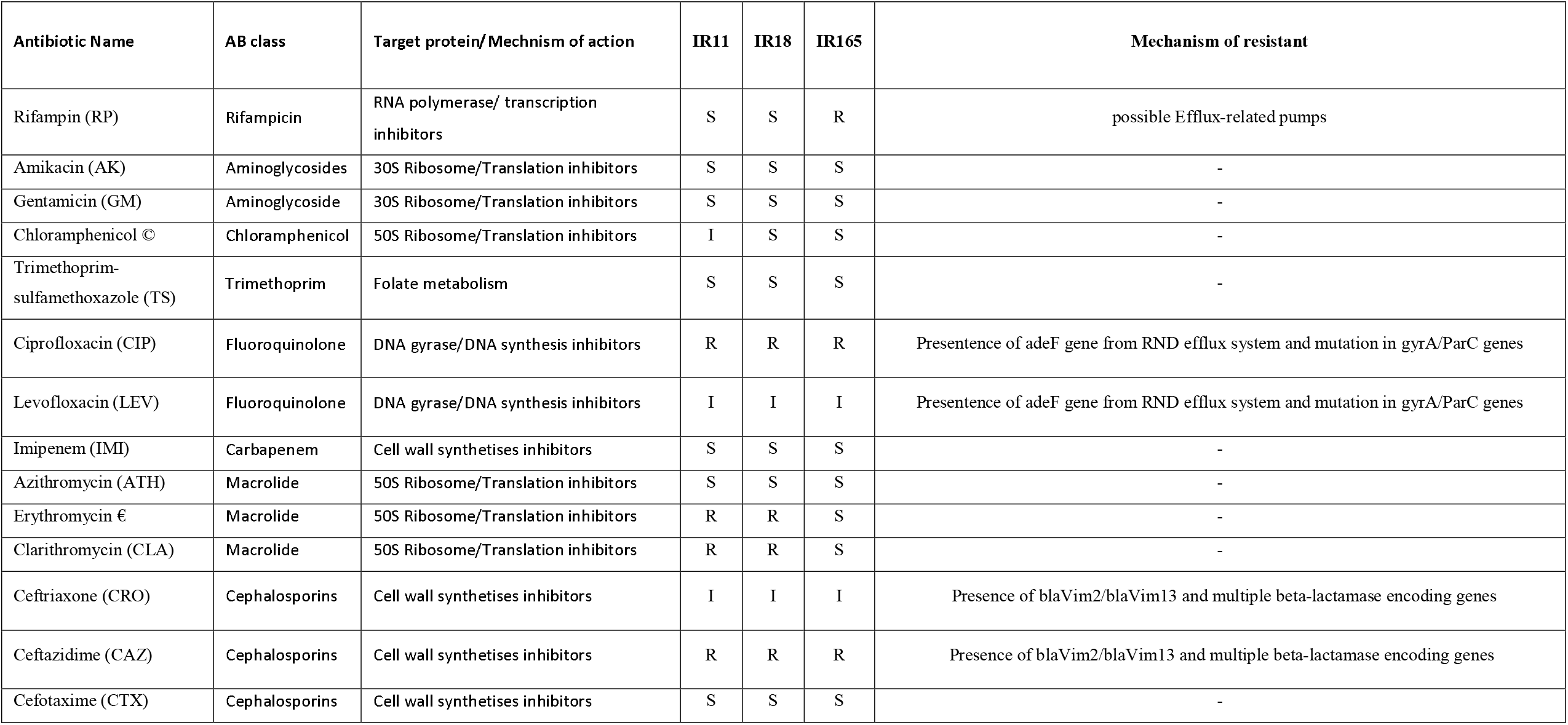
Antibiotic resistant test results and resistance mechanisms of clinical isolates in this study

Genome investigation and genome annotation analysis of two isolates using CARD [24] revealed the presence of *adeF* gene and other efflux pump genes that are involved in the antibiotic transportation systems in the genomes (Supplementary file-2). Mutations in the *gyrA* (Ala-83 and Met-87) and *parC* (Gln-57, Val-66 and Ala-80) encoding DNA gyrase and Type IV topoisomerase, respectively were detected.

IR11 and IR18 strains were found to be resistant to macrolide antibiotics, erythromycin and clarithromycin, possibly due to the presence of macrolide-specific efflux protein MacA and Macrolide-specific *ABC*-type efflux carrier (TC 3.A.1.122.1) (supplementary file-2) that were absent in IR165. In contrast, IR165 was resistant to rifampin due to possible efflux-related mechanisms that pump out the antibiotics from the cells.

### 3.6 Comparative genomic analysis of *B. diminuta* isolates

A total of 11 available sequenced genomes of *B. diminuta* (as of September 2020) with an average size of 3.36 Mbp and 3212 average protein coding regions. These isolates were collected from various environments such as water, gentamicin fermentation residue, and cheese. Four isolates collected from human causing co-infections along with other pathogens (Table 3). Pan-genome analysis of the 13 genomes (11 global and two Iranian isolates) using Roary (v3.12.0)[22] revealed that from 8233 genes as pan-genes, the core genome consists of 1501 genes (on average 45.65% of each genome [1501/3288]), with the accessory genomes containing 3407 genes in the shell (41.38% of pan-genome). The unique genes ranged from 7 for ACCC1057 to 1105 for 470-4 (Fig-2. A). We analysed the pangenome of the six isolates that were collected from humans showing 1590 core genes from the total of 6512 genes. We found no specific common genes for these genomes when compared to the environmental-collected isolates (Fig-2B).

**Figure-2.A.**
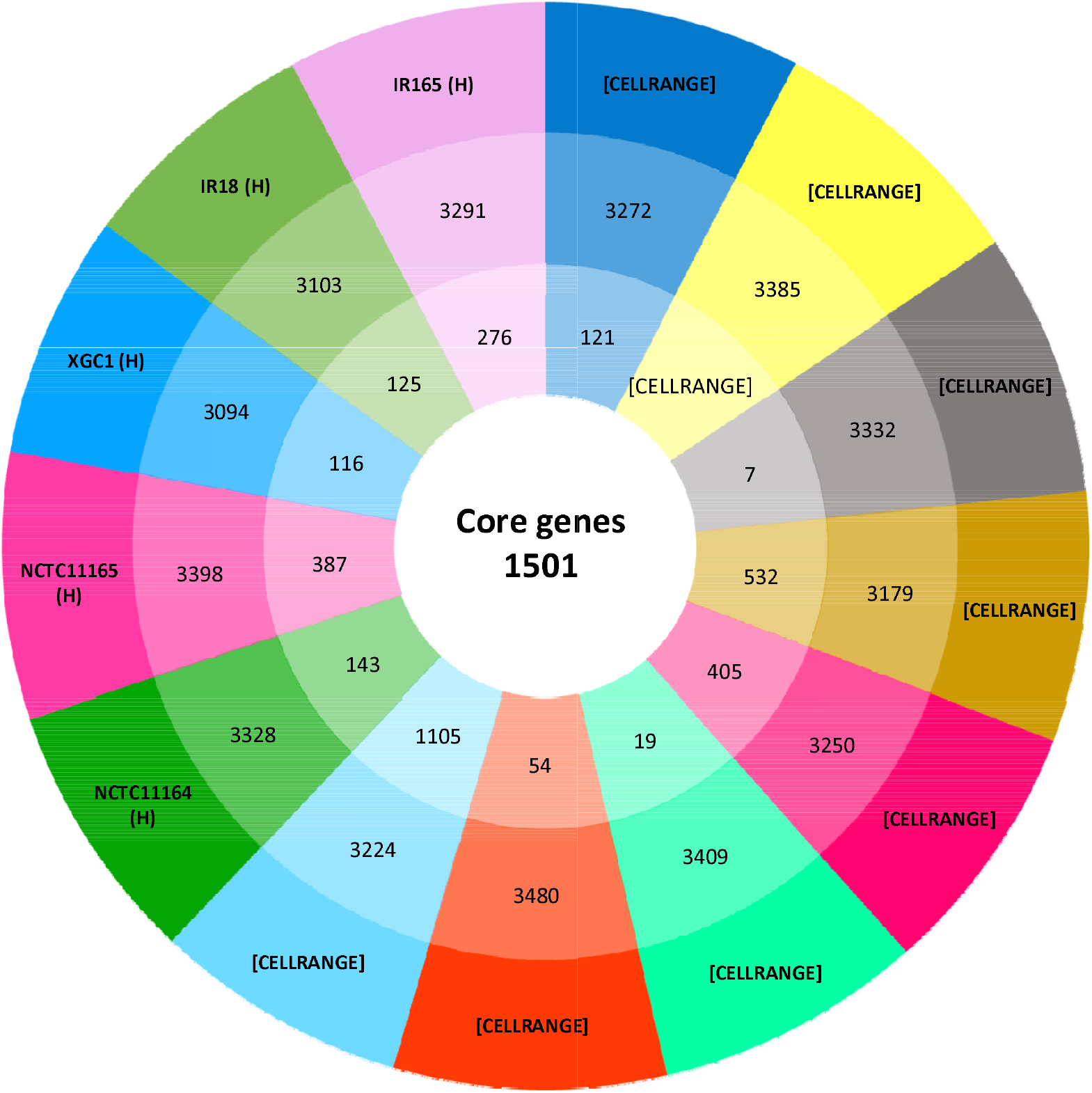
Circular genome representation of 13 *B. diminuta* isolates collected worldwide. The intersection of all strains presents the total number of core genomes. The intersection of each pair represents the total number strains, while the outer number represents the total number of genes and the inner Number represent the unique genes.

**Figure-2.B:**
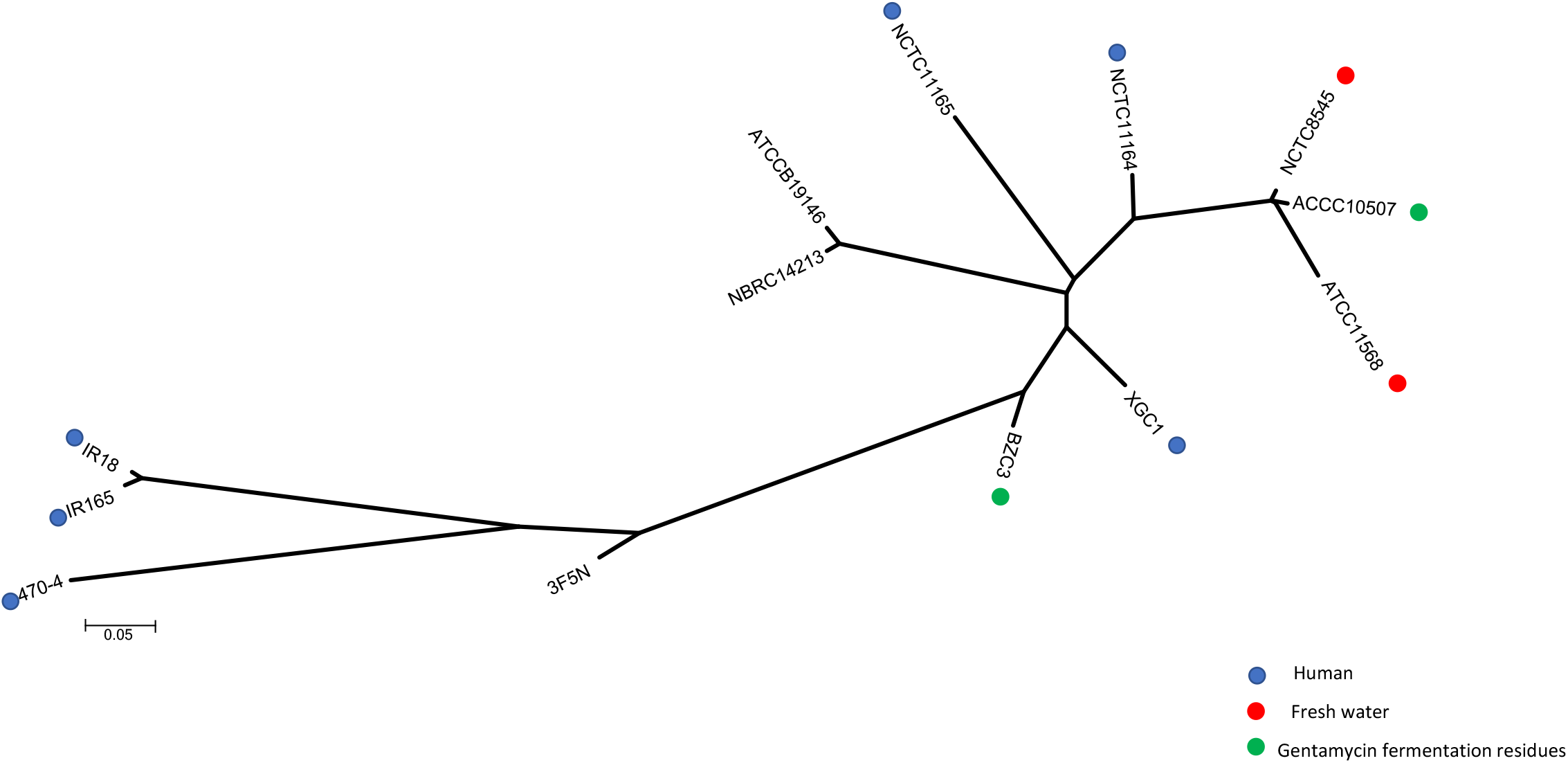
Phylogeny analysis of 13 Iranian and global *B. diminuta* isolates using core genome (by Prokka and Roary) for maximum parsimony analysis (Mega7). The Iranian isolates grouped with (470-4), the clinical oral sample isolated from the patient in USA

**Table 3-.** *B. diminuta* isolates used for global phylogeny analysis and their genomic characterizations.

| strains | size<br>(Mbase<br>) | GC<br>% | No.<br>genes | No.<br>proteins | rRNA | tRNA | Pseudogenes | Year of<br>submission | Country | Source of Isolation | Seq. method | Biosample<br>submission Number |
| --- | --- | --- | --- | --- | --- | --- | --- | --- | --- | --- | --- | --- |
| ATCC11568 | 3.37 | 66.3 | 3398 | 3151 | 3 | 47 | 193 | 2011 | USA | Freshwater water | 454 GS FLX | SAMN02469913 |
| NCTC8545 | 3.44 | 67.3 | 3425 | 3333 | 6 | 51 | 32 | 2014 | UK | Freshwater water | Pac Bio | SAMEA4040024 |
| ACCC10507 | 3.4 | 67.3 | 3406 | 3302 | 4 | 50 | 46 | 2017 | China | Gentamicin fermentation<br>residues | Illumina hi seq | SAMN07236627 |
| 3F5N | 3.25 | 67 | 3233 | 3146 | 3 | 47 | 33 | 2017 | France | Soft smear-ripened cheese<br>with raw milk | Shotgun<br>sequencing | SAMEA80388418 |
| BZC3 | 3.39 | 67.4 | 3294 | 3197 | 6 | 52 | 35 | 2017 | China | Gentamicin fermentation<br>residues | Illumina hi seq | SAMN07236578 |
| NBRC14213 | 3.5 | 67.5 | 3461 | 3351 | 3 | 46 | 57 | 2019 | Japan | A contaminant in a culture<br>of *Bacillus cereus* | Shotgun<br>sequencing | SAMD00097246 |
| ATCCB1914<br>6 | 3.55 | 67.4 | 3506 | 3391 | 6 | 51 | 54 | 2019 | China | unknown | Oxford<br>Nanopore | SAMN10737780 |
| 470-4 | 3.34 | 66.9 | 3298 | 3136 | 3 | 45 | 111 | 2012 | USA | Oral cavity | Illumina | SAMN02436690 |
| NCTC11164 | 3.41 | 67.4 | 3383 | 3274 | 6 | 51 | 48 | 2017 | UK | Human, cerebrospinal fluid | PacBio RS | SAMEA104338351 |
| NCTC11165 | 3.35 | 67.3 | 3332 | 2981 | 9 | 54 | 284 | 2017 | UK | Human, throat swab | Shotgun<br>sequencing | SAMEA104167223 |
| XGC1 | 3.18 | 67.4 | 3170 | 3075 | 4 | 45 | 42 | 2018 | India | Tuberculosis Patient-blood | Illumina<br>NextSeq | SAMN03480409 |
| IR18 | 3.16 | 67.2 | 3221 | 3100 | 3 | 49 |  | 2020 | Iran | Nasopharynx | Illumina | SAMN17214354 |
| IR165 | 3.34 | 67.1 | 3419 | 3368 | 3 | 50 |  | 2020 | Iran | Nasopharynx | Illumina | SAMN17214355 |

The maximum parsimony analysis showed that isolates collected from human specimens were dispersed in the tree and grouped with the ones collected from other sources including water, soil, or gentamicin fermentation residue. Two Iranian isolates were grouped in one branch together with *B. diminuta* 470-4 which was isolated from the oral cavity in the USA and submitted in NCBI in 2012 (Supplementary Figure1).

## 4. Discussion

*B. diminuta* is commonly used for testing of sterilizing-grade filter retention due to its characteristics and has been defined as an opportunistic pathogen causing infections in different part of the body with a rare report of respiratory tract infection [7, 8, 12].

Infection caused by *B. diminuta* has been reported from patients with an underlying conditions or diseases such as cancer or urinary tract infections that allowed patients to succumb to *Brevundimonas* infections [4]. Isolation of *B. diminuta* from two patients with pertussis-like symptoms was previously reported from Pasteur Institute of Iran [23]. Here we isolated three additional *B. diminuta* strains from patients with respiratory tract infections with symptoms like vomiting, severe cough and fever (Table 1). Interestingly two isolates, IR11 and IR18, were from infants under six-month-old age which was not commonly reported previously.

*B. diminuta* and *B. pertussis* have some similar biochemical and morphological similarities (Appendix-1) including Gram stain, colony morphology, and positive catalase and oxidase results. Therefore, it might be difficult to differentiate *B. pertussis* and *B. diminuta* at the first step in diagnostic laboratories based on these tests alone. This may lead to false pertussis reports if there are respiratory tract infections especially when there are respiratory symptoms in infants. In this case, it is suggested to perform some additional molecular tests including real-time PCR targeting *ptxP*, the promoter of pertussis toxin, IS*481* and IS*1002* which are two insertion sequence in *B. pertussis* [26]. According to new pertussis guidelines and policies implemented in 2009 by Iran Centre for Communicable Disease Control [25] all nationwide suspected samples from patients with pertussis symptoms must be notified. Furthermore, samples including swabs and cultures must be sent to the reference laboratory of Pasture Institute of Iran to confirm disease. For these three cases swabs were sent to the reference laboratory for pertussis confirmation, the real time PCR results for IS*481*, IS*1002* and *ptxP*, were negative. Therefore, an additional biochemical characterization test (API E20) and 16s rRNA sequencing were performed and the results confirmed them to be *B. diminuta*. There is little information about pathogenicity or mechanisms of resistance in *B. diminuta*. Antimicrobial resistance was reported for *B. diminuta* globally especially for fluoroquinolones, β-lactams, aminoglycoside and colistin [5, 7, 27]. All three clinical isolates in this study showed resistance to ceftazidime and intermediate resistance to levofloxacin. Ciprofloxacin and levofloxacin belong to the fluoroquinolone class of antibiotics that target DNA synthesis and successfully used against a wide range of gram-positive and gram-negative bacteria. Fluoroquinolones bind to DNA gyrase or topoisomerase proteins then affect the DNA synthesis in bacteria [37]. Different mechanisms are reported to be associated with the bacterial resistance to ciprofloxacin including the presence of proteins known as efflux pumps or mutation in quinolone-target proteins and efflux pumps [38]. The genomic investigation using antibiotic resistance databases revealed the presence of *adeF* gene in the genomes leading to the pumping fluoroquinolones out of the cell and resulting a high level of resistance. AdeF protein is the membrane fusion protein of the multidrug efflux complex AdeFGH pump that is involved in the transportation of fluoroquinolone antibiotics out of the cell. AdeFGH pump belongs to Resistance-nodulation-division (RND) superfamily efflux pumps which is one of the most important systems in mediating antibiotic resistance in Gram-negative bacteria [39]. The RND systems have various functions, including extrusion of toxic compounds, homeostasis of the cell, and export of virulence determinants [40]. They generally exhibit a broad substrate spectrum, and their overexpression can diminish the activity of several unrelated drug classes [41].

Another mechanism that resulted resistance to the fluoroquinolone family is the mutation in QRDR (the quinolone resistance-determining region) of *gyrA* and *gyrB, parC and paeE* gene of which mutations in gyrA and parC are more frequent [5, 42, 43]. The most-reported mutations in gyrA, encoding subunit A DNA gyrase, are found in the amino acid at positions 83 (*Ser*/*Thr*-83) and 87 (*Asp*/*Glu*-87) of the QRDR *GyrA* subunit for quinolone resistance mechanism in pseudomonas [5, 38]. The *GyrA* QRDR of all available *B. diminuta* isolates were compared with *P. aeruginosa* and *B. diminuta* ATCC 11568, isolated in 1952 [44]. All *B. diminuta* isolates have *Ala*-83(92 in our genomes) and *Met*-87(96 in our genomes) mutation in the *GyrA* QRDR region, confirming the intrinsic quinolone resistance as previously reported for clinical isolates [5, 6].

Furthermore, the mutation in type IV topoisomerase, as another target protein for ciprofloxacin, is another mechanism that is believed to be responsible for quinolone resistance [37, 38]. The enzyme is a heterotetramer protein encoded by *parC* and *parE* genes and a mutation in QRDR of *ParC* is more frequent and effective in ciprofloxacin resistance bacteria including *P. aeroginosa* [37]. The mutations in *parC* gene in *B. diminuta, Gln*-65, *Val*-74, *Ala*-88 and *Glu-92* were identified in all Iranian and global genomes as the third mechanism for fluoroquinolone resistance as reported in *B. diminota* ATCC 11568 and other clinical isolates [5].

The isolates in this study shown also resistant to ceftazidime and intermediate resistance to ceftriaxone. Both antibiotics are the third generation of cephalosporins which are categorised as the β-lactam class of antimicrobials that target bacterial cell wall proteins known as Penicillin Binding Proteins (PbPs) [45, 46]. Multiple mechanicians are associated with resistance to cephalosporines mostly dependent on β-lactamase proteins such as IMP and VIM [47, 48, 49].

Here, we detected *blaVIM-2* and *blaVIM-13* genes along with other β-lactamases-mediated genes in isolated genomes which encode metallo-β-lactamase proteins that hydrolyse cephalosporins. These genes were also detected in different bacteria including *P*.*aeruginosa* as well as clinical and environmental *B. diminuta* isolates [50, 51, 52].

Macrolide including erythromycin and clarithromycin has a broad spectrum of antimicrobial activity on the bacterial ribosomal RNA (rRNA) subunit [53, 54]. These drugs target bacterial protein synthesis by inhibiting the extension of the peptide chain [53, 54, 55]. Modification of macrolide-binding site in the 23S ribosomal RNA (rRNA), demethylation, methylation or mutation within the 50S ribosomal subunit, efflux pump systems and ATP-binding cassette (ABC) family of transporters are some of the known macrolide-resistance mechanisms in bacteria [56, 57]. Here, we found two macrolide-specific efflux pump genes including *macA*, Macrolide-specific ABC-type efflux carrier (TC 3.A.1.122.1), ABC-type multidrug transport system and ATPase component in IR18. These genes encode proteins that are shown to mediate the high level of macrolide resistance by transporting the antibiotic outside the cell [58].

Rifampin is a synthetic derivative of natural products of the bacterium, with an inhibitory effect on beta-subunit of the RNA polymerase of prokaryotes and prescribed for gram-positive and gram-negative bacteria [59, 60]. Resistance to rifampicin (RIF) is mediated by mutations clustered in a small region of the *rpoB* gene [61, 62] which encodes the RNA polymerase β-subunit, or possibly by multidrug efflux pumps [59]. Previous studies confirmed mutation in the different amino acid sequences of *rpoB* protein mainly in 516, 526 and 536 mediated rifampin resistance in bacteria [60]. However, we found no mutation in these positions in the *rpoB* gene of IR165 and the alignment investigation shown 100% similarity of the amino acid sequence of IR165 with IR18 as rifampin-sensitive isolates. Therefore, resistance to rifampin in isolate IR165 might be due to the efflux-related mechanisms as reported previously [63] and needs further investigation.

Currently, whole-genome sequencing has largely been used as a research tool to expand our knowledge about microbial genomic characteristics and large-scale genome analysis, epidemiological investigations, and evolutionary studies. To our knowledge, there is no study to investigate the relationship between *B. diminuta* isolates or to define the core genome throughout them. Core genome alignments could be used as a suitable approach for phylogeny analysis and modelling genomic diversity. Across the full set of 13 *B. diminuta* including Iranian isolates, the average genome has less than 50% core genes (1501 gene/average genome size 3288[45.65%]) as genes that are present in all genomes which is in line with other bacterial species with an average core genome of 1650 for bacterial species showed in previous studies [64]. However, pan-genome analysis depends on various factors such as sequencing platforms, the percentage identity cut-offs and the number of genomes included [64]. It also depends on whether the species has an open pan-genome as an increasing pangenome when a new genome is added [64, 65]. The pan-genome size of these isolates was 8233 and their increased tendency showed it can be taken as an open pan-genome due to the vast diversity among the isolates (Fig.3). The high number of cloud genes (3325 [40.38% of pan-genome]) as the number of genes in less than 15% of the genomes may help the bacteria for specific environmental niche adaptation due to the source of isolation that may influence the bacterial adaptation. since most of the available isolates were collected from non-clinical environments (7/11[63%]) such as water or fermentation residue.

**Figure-3:**
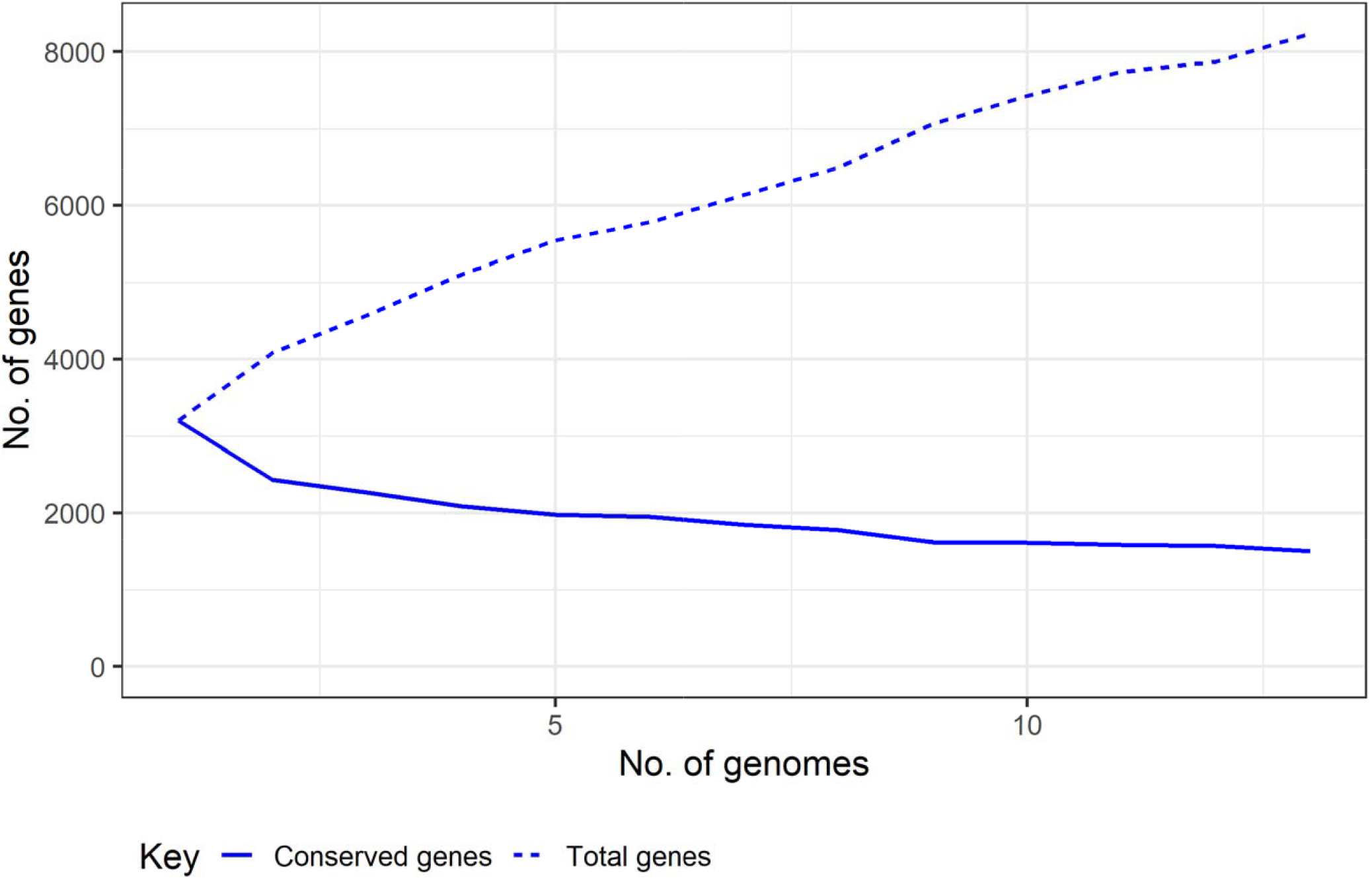
Heap’s law chart representation regarding conserved genes and total genes in 13 *B. diminuta* genomes showing an open pan genome for *B. diminuta* reflecting diversity of isolation sources.

## 5. Conclusion

*B. diminuta* is currently considered as an opportunistic pathogen. However, in recent years it is reported as causative agent of different infections in humans such as respiratory tract infections. In this study, we isolated *B. diminuta* from patients mostly children under five years old who initially diagnosed with whooping cough caused by *B. pertussis*. Our findings including bacterial characterization, 16s RNA analysis, whole genome sequencing and antimicrobial susceptibility test showed these isolates are *B. diminuta*. The three isolates have a different level of antimicrobial resistance to quinolone and cephalosporine with multiple mechanisms including some intrinsic mechanisms. We also detected core and pan genes of available global *B. diminuta* and perform a phylogenetic analysis that showed our isolates grouped with one clinical *B. diminuta* collected from the USA. These findings have important implications on the differentiation of infections caused by *B. diminuta* especially for hospitalised respiratory patients and antibiotic treatment policies for patients.

## Acknowledgments

This work was supported financially by Pasteur Institute of Iran.

## Figure legends

**Supplementary Figure-1:**
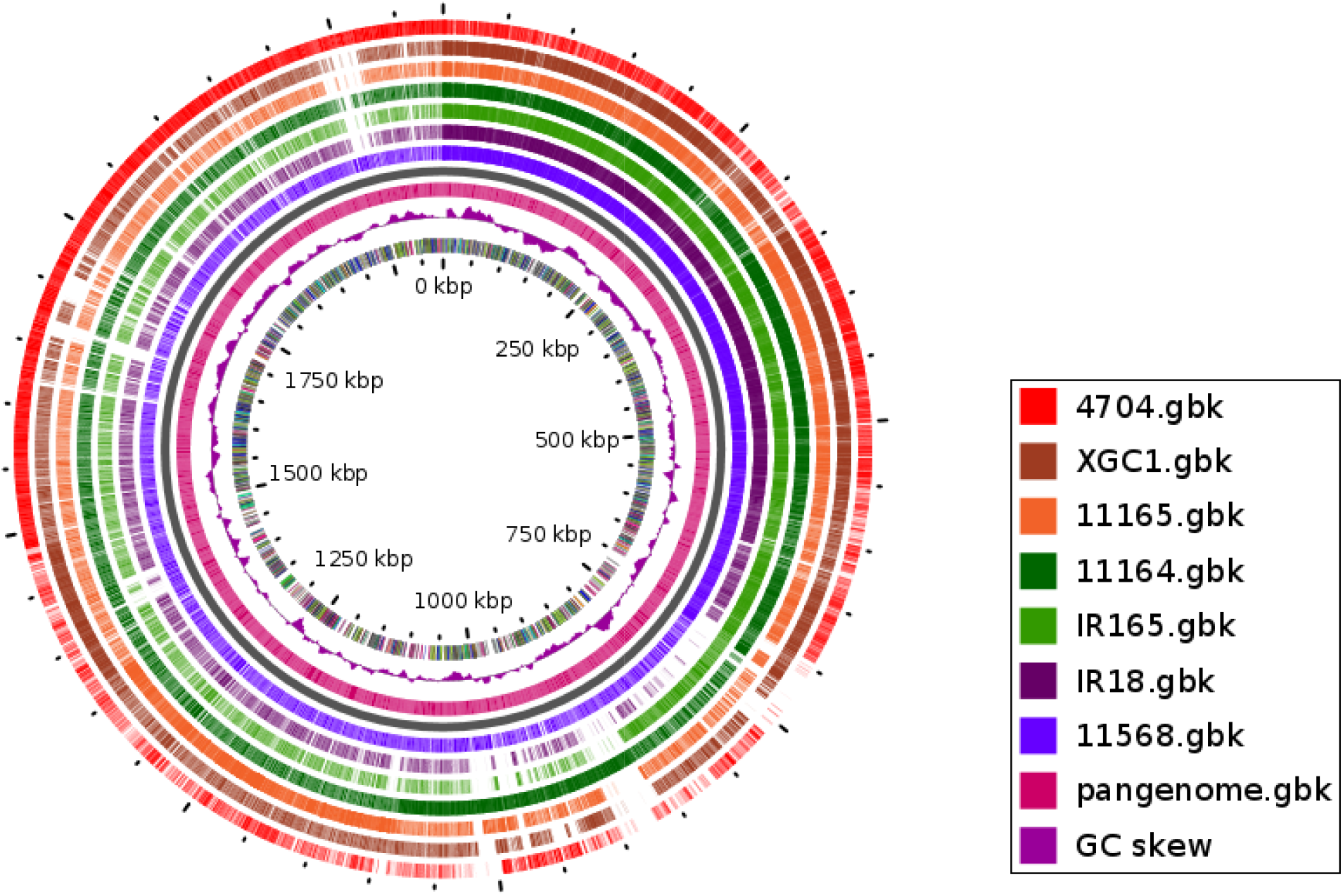
BLAST Atlas comparing six clinical *B. diminuta* isolates from different countries against *B. diminuta* strain ATCC11568 as a reference genome. Circular plot generated using G view Comparison Tool (https://server.gview.ca/) using BLASTn, G + C skew (purple)

## Tables

**Appendix-1:** Bacterial culture and Biochemical characteristics of *B. pertussis* and *B. diminuta*

| Basic Characteristics | *Bordetella pertussis* | *Brevundimonas diminuta* |
| --- | --- | --- |
| Gram Staining | Gram-negative (-ve) | Gram-negative (-ve) |
| Catalase | Positive (+ve) | Positive (+ve) |
| Oxidase | Positive(+ve) | Positive(+ve) |
| Urease | Negative (-ve) | Negative (-ve) |
| Nitrate Reduction | Negative (-ve) | Negative (-ve) |
| Gelatin Hydrolysis | Negative (-ve) | Negative (-ve) |
| Growth in nutrient broth | Negative (-ve) | Positive (+ve) (poor and slow growth) |
| Growth on MacConkry | Negative (-ve) | Negative (-ve) or poor and slow growth |
| Haemolysis | Positive (+ve) | Negative (-ve) |
| Flagella | Negative (-ve) | Positive (+ve) |
| Motility | Negative (-ve) | Positive (+ve) |
| Pigment | Negative (-ve) | Positive(+ve) |
| Shape | coccobacilli (4x8µm) | coccobacilli (4x5µm) |
| Spore | Negative (-ve) | Negative (-ve) |

